# Sustainability During Instability: Long-Lived Life Science Databases and Science Funding Outlook in the United States

**DOI:** 10.1101/2025.10.08.680785

**Authors:** Heidi J. Imker

**Affiliations:** University Library, University of Illinois Urbana-Champaign, Urbana, Illinois 61821

## Abstract

For decades, life science researchers have had cost-free, unrestricted access to data through online databases. However, the sustainability of even well-established resources was already tenuous, and abrupt changes in science funding in the United States seems poised to exacerbate these challenges. This study employed a multiple mini case study approach, triangulating semi-structured interviews with supplemental documentation to investigate 9 diverse, long-standing databases that had previously received support from U.S. federal agencies. The research explored each database’s purpose and use, examined current and emerging funding strategies, and considered the potential consequences if any of these databases were forced to shut down. Participating databases include: 1) BHL: Biodiversity Heritage Library, 2) MorphoBank, 3) OMIM: Online Mendelian Inheritance in Man, 4) ORDB: Olfactory Receptor Database, 5) rrnDB: ribosomal RNA operon copy number database, 6) VEuPathDB: Eukaryotic Pathogen, Vector, and Host Informatics Resources, 7) WormAtlas, and 8-9) two databases that wished to remain anonymous. Findings revealed anticipation of higher barriers to data access and reuse, loss of subject matter expertise now and into the future, and lost or interrupted opportunities. Additionally, real impacts have already begun through redirection of energy, abrupt reductions in support, and increased competition for funding. Sustainability models in light of the current funding outlook in the US are discussed.

## Introduction

Cost-free, unrestricted access to collections of data has been a hallmark of life science research for decades. These collections, often in the form of open, online databases, also referred to more broadly as “biodata resources”, are extraordinarily diverse, covering a wide range of data, topics, and methods. They range in scale from well-established “core” resources that support wide swaths of the life sciences community to more focused databases dedicated to specific research areas, such as a distinct methodology, disease, or organism. Each aims to provide a user community, large or small, with access to high-quality data, with deep expertise informing data curation, standardization, metadata application, and resource design. Ultimately, these resources represent focused collections of Subject Matter Expertise (SME), applied to life sciences data, that enable discovery, reference, computation, de novo analysis, and metanalysis. Additionally, these resources also often include educational and training materials as well as analysis and workflow enhancement tools, all aimed at further advancing knowledge in the field.

What is otherwise a tremendous strength has a downside. Sustaining these resources has been a long-standing challenge and the focus of numerous reports and studies (Chandras et al. 2009; Crow 2013; Jarvenpaa and Essén 2023; OECD 2017; Schofield et al. 2010; Shankar 2016; Strecker et al. 2023). Biodata resources are nearly always launched through short-term funding mechanisms—typically via project grants or contracts, with commensurate closeouts and termination dates, rather than as infrastructure intended to serve for years into the future. While not every data resource ought to be permanent, there are databases that have proved their utility through decades of existence and thousands of users. However, periodic changes in funding priorities, like the current shifts at major U.S. federal agencies like the National Institutes of Health (NIH) and the National Science Foundation (NSF), compromise resource stability and the science that data resources enable.

Starting the recent shifts, tracing funding cuts through time demonstrates the persistence of the challenge. In September of 2024, NIH’s National Institute of Allergy and Infectious Diseases (NIAID) cut all funding to VEuPathDB (a database represented in this study), which provides a suite of databases used to study pathogenic organisms (Wadman 2024). A few years earlier, 2021 saw continued concern for model organism databases after suffering years of significant cuts following the controversial closure of the NIH’s National Center for Research Resources (Bellen et al. 2021; Oliver et al. 2016; Wadman 2010). Farther back, The Arabidopsis Information Resource began charging users in 2013 after the NSF began phasing out funding in 2009 (Abbott 2009; Check Hayden 2013). Similarly, a 2007 decision by NIH’s National Library of Medicine to stop funding infrastructure in favor of innovation led to a series of cuts for 5 databases (Baker 2012). Nor is the issue limited to the 21st century. *Nature* reported on funding issues for several prominent databases in 2005, with a representative for the Mouse Genome Database noting the challenge was already a decade in the making (Merali and Giles 2005). In 1998 Ellis and Doyeb published in *Nature Biotechnology* on the potential for data loss, with a pointed abstract which reads in its entirety “Data rich but cash poor, many free biological databases are on the verge of financial collapse.” (Ellis and Kalumbi 1998). That article reported that of the nearly 400 databases examined, two-thirds were uncertain of future funding. In fact, while database proliferation did accelerate in the 1990s, the question of sustainable funding has dogged the life sciences from the very beginning. Over 40 years ago, licensing intended to generate a windfall in case of funding loss played a role in determining who won the grant contract for the first version of GenBank (Strasser 2019). Prior to the digital age, and concurrently into the present, sustaining physical data collections used for natural history and comparative biology has likewise suffered on-going challenges (Miller et al. 2020).

The US has made outsized impacts on life sciences research worldwide by having a substantial financial and intellectual stake in establishing and maintaining online biodata resources (Imker 2020). However, throughout this time, the need for a cohesive, sustainable strategy has been repeatedly recognized by the research community and US funding agencies (*Biological Collections* 2020; Bourne et al. 2015; National Science Board 2005). The international scope is also a challenge, since data is often sourced from throughout the world and the user communities are global, as well. The Global Biodata Coalition (GBC) was established as a forum for research funders across the world to share knowledge and develop strategies to support biodata resources (Anderson 2017). In the time since its launch, GBC has worked to raise awareness, identify open access core resources that support large portions of life science research, describe the biodata infrastructure landscape across the world, and encourage conditions that enable funders to coordinate across international borders (Imker et al. 2023). As of 2025, two US federal funding agencies, NSF and NIH, are among its international membership.

Despite awareness and acknowledgment of the challenge, the problem persists today, and the sudden disruptions to science funding in the US in early 2025 signaled the potential for a fresh wave of crises for life science data resources, particularly those that relied on federal funding in the past. Often concern arises database by database, so to provide a wider view, this study profiles 9 long-lived, online life science databases—each varying in data type, scope, and scale—to capture today’s landscape. It highlights common challenges and projected impacts, aiming to inform efforts to strengthen the broader ecosystem.

## Methods

### Methods Overview

To examine new federal funding cuts as an emerging phenomenon, this study employs an exploratory, interpretive multiple mini case study (MMCS) approach with an emphasis on cross-case analysis (Käss et al. 2024; Yin 2018). The principal question was to investigate if long-lived life-science data resources are particularly vulnerable amid wide-spread cuts to science funding and to document their challenges at this moment in time. The study proposal was reviewed by the University of Illinois Urbana-Champaign Office for Protection of Research Subjects and designated as minimal risk (IRB25-0306). All activities were conducted solely by the author using video, data storage, and analysis platforms licensed for use by the University of Illinois Urbana-Champaign.

### Database Selection and Primary Source Material Collected

Individual databases served as the unit of analysis and were selected using the following criteria to address the scope and purpose of the research question: each must 1) be based in the US with a focus on life science data relevant to academic research, 2) have existed freely available via world wide web access for a minimum of 15 years, not including any prior years when the data collections may have been available other online mechanisms, such as dialup, or distributed within other collections, and 3) have sufficient public documentation, either in articles, the database website itself, or other sources, to triangulate details such as database purpose and having received federal funding from the US government. Beyond these criteria, databases were purposely targeted to represent the wide diversity of resources that make up the larger ecosystem, including a variety of types of data (e.g., image, sequence, etc.) as well as a variety of scales and scopes (e.g., focused on a specific protein class versus a broader topic). Interviews of database representatives, identified as a corresponding author on associated database articles or publicly named as contacts on database websites, formed the anchor of each case. Additional primary sources were collected, including live database webpages, archived WayBack Machine webpages, academic articles, and other documentation such as published reports and news stories (Table S1. Primary Source Material Counts per Database, Table S2. Source Material Profiles).

### Protocol

Interviews were conducted to gather first-hand information on each database’s purpose, origin, and current funding, as well as to understand the potential impact of funding cuts. Prior to each interview, primary sources for each database were collected and reviewed with details noted in a pre-interview document and then referred to during the interview, as appropriate (e.g., verification or clarification of current funding status). Semi-structured interviews with open-ended questions were chosen to allow participants to take the conversation in directions that seemed most relevant to them and their database’s circumstances (Table S3. Interview Prompts). Representatives from 15 long-lived online databases were invited to participate via email, with interviews covering 9 databases (10 participants total) held in the summer of 2025. A 10th resource initially accepted the invitation but then withdrew prior to the interview due to on-going uncertainty. Participants had the option of being explicitly named, along with the identity of their database, or remaining anonymous. Of the 9 databases represented, 2 choose to remain anonymous. For these resources, referred to throughout as DB1 and DB2, care was taken to ensure that no specific information about the representative or the resource (e.g., specific focus, age, location, type of data collected, or funding) was revealed. Interviews were conducted using either Zoom or Microsoft Teams, with transcription and recording of the video and audio when approved by the participant. When recording was not agreed to, hand-typed field notes were carefully reviewed immediately following the interview to edit for accuracy; when recording was approved, transcripts were checked against the video and audio and annotated to correct where needed. Following interviews, additional primary source materials were collected as needed to improve the accuracy of the author’s interpretation.

### Coding, Analysis, and Reporting

All primary data sources were collected in a case study database and analyzed using MAXQDA 24 software. Primary sources, interview recordings, and notes were reviewed and iteratively coded, cross checking between sources throughout, following the conventional content analysis approach (Deterding and Waters 2021; Hsieh and Shannon 2005; Timmermans and Tavory 2012). Higher-level themes were developed through multiple, successive interactive reviews of analytic codes (Ryan and Bernard 2003). Visualization tools in MAXQDA were used to identify analytical codes that occurred across the majority of the database, suggesting commonalities; these were then repeatedly reviewed reflectively and analyzed further to verify and synthesize observations (see Supplemental Codebook for the codes, definitions, and examples, Table S4. US Federal Agency Abbreviations, and Figure S1. Code Matrix). Last, specific observations or examples from participants that illustrated themes were noted in analytical memos to be described in more detail or quoted directly. Quotes were edited to remove filler phrases (“you know”) and vocalizations (“um”) to avoid detracting from the point of participants’ statements. As an additional measure of accuracy, participants were sent a draft of the manuscript and asked to review the information associated with their specific database, with 8/9 contributing.

## Results

### Overview of Database Cases

Databases are distributed across the US and launched on the web between 18 to 31 years ago. Brief database profiles are provided in Table 1 with more comprehensive overviews in Table S5.

**Table 1.** Database Profiles in Brief

| **Name**<br>URL, 1<sup>st</sup> Year on Web, Current Host | **Primary Data** | **Primary Uses** |
| --- | --- | --- |
| **BHL: Biodiversity Heritage Library**<br>• biodiversitylibrary.org<br>• 2007<br>• In transition from the Smithsonian Institution | Data and metadata from high resolution archival (scans) and contemporary (born digital) literature and archival materials of biodiversity relevance | Used as primary source material to study historical trends in biodiversity, verification of taxonomic names and classifications; Used for educational and artistic purposes |
| **DB1: Database 1**<br>• > 15 yrs ago<br>• US Organization | Data of biomedical relevance | Used to access high-quality data and tools relevant to the study of human health |
| **DB2: Database 2**<br>• > 15 yrs ago<br>• US Organization | Data of biomedical relevance | Used to access high-quality data and tools relevant to the study of human health |
| **MorphoBank**<br>• morphobank.org<br>• 2004<br>• Phoenix Bioinformatics | Phylogenetic matrices and associated data (taxa, characters and states, cells with or without images, bibliographic citations), 2D and 3D media of specimen morphology | Used to build on previous research and support large-scale comparative studies |
| **OMIM: Online Mendelian Inheritance in Man**<br>• omim.org<br>• 1995<br>• Johns Hopkins University | Gene and phenotype entries with relationship mapping, clinical summaries, and references to corresponding literature and external data sources | Used as the “first stop” authoritative resource for exploration of genes and genetic disorders; Used in medical education at multiple levels |
| **ORDB: Olfactory Receptor Database**<br>• ordb.biotech.ttu.edu/ORDB<br>• 1994<br>• Texas Tech University | Nucleotide and amino acid sequences for olfactory receptor (OR) and olfactory receptor-like (ORL) gene and protein sequences with interoperability between research labs and partner databases | Used to study OR and ORL genes, access to keys for variable lab IDs, and access tools to analyze unpublished sequences |
| **rrnDB: ribosomal RNA operon copy number database**<br>• rrndb.umms.med.umich.edu<br>• 2001<br>• University of Michigan | 16S and 23S RNA genes (sequencing and earlier experimental), organism taxonomies, statistical summaries, additional metadata and links to related external resources | Used for basic research (initial) with subsequent practical use to counteract bias while measuring composition of microbial populations |
| **VEuPathDB: Eukaryotic Pathogen, Vector, and Host Informatics Resources**<br>• veupathdb.org<br>• 2004<br>• Universities of Pennsylvania and Georgia | Genome sequence and annotation, transcriptomics, proteomics, epigenomics, metabolomics, population resequencing, clinical data, surveillance data, host-pathogen interactions, and orthology profiles | Used for centralized access to multi-omic data and associated specialized tools to explore, analyze, and compare genomic and functional data on pathogens, vectors, hosts, and disease |
| **WormAtlas**<br>• wormatlas.org<br>• 2002<br>• University of Illinois Urbana-Champaign | Expertly annotated electron microscopy images organized by species, tissue, genotype, age, or sex with corresponding detailed anatomical maps, handbooks, and interactive educational resources | Used to compare and study anatomical features, cellular structures, identification of cellular location of gene expression, developmental processes, and phenotype variations; Used for educational materials and in teaching |

Each resource has a distinct focus and purpose, as reflected in the data collected and the expertise applied, and can broadly categorized with the following foci: 1 model organism (WormAtlas), 1 protein class (ORDB), 1 RNA class (rrnDB), 2 biodiversity (BHL and MorphoBank), and 4 on human health conditions (OMIM, VEuPathDB, DB1, and DB2). They range widely in the scale of their operations, which is apparent in the scope of the resource, features and complexity in their systems, collection size and growth over time, and the tools and educational materials developed—these all reflecting the respective financial, technical, and human resources available to each (Table S5. Database Profiles).

Collectively, the audiences cover a wide range of the life sciences, including: bioinformaticians, botanists, computational biologists, cell biologists, developmental biologists, ecologists, epidemiologists, evolutionary biologists, geneticists, microbiologists, mycologists, neuroscientists, paleontologists, parasitologists, systemists, taxonomists, and vector biologists, plus those in other disciplines such as computer science and history. Audiences also extend beyond academia to include professionals in the medical field (such as physicians, clinicians, and public health experts) as well as individuals in education from high school students and teachers to postgraduate learners and educators. BHL, in particular, has a robust public audience of artists and hobbyists.

### From Inspiration to Implementation

While each database serves a distinct purpose, the impetus for all 9 resources was rooted in the same high-level need: efficient use of prior science by making otherwise scattered or inaccessible data discoverable and usable (Table S3. Database Profiles). Each accomplishes its goal by using SME, referred throughout as just “expertise” to carry out functions such as identifying and aggregating relevant data, curating for quality and standardization, organizing, and enriching with the contextual information needed in the field of study. Prior to resource launch, documentation and interviews cited issues with researchers having to personally track down, evaluate, extract, and decipher data. When possible at all, this was reported as time consuming, inconsistent in terms of access, and left the data susceptible to varying interpretations. The value of providing authoritative, expertly curated data to address confusion or disagreements in the field was expressed by 7 database representees (DB1, DB2, MorphoBank, OMIM, ORDB, rrnDB, and WormAtlas).

In several cases, the database provides access to data otherwise not available at all. DB1, DB2, MorphoBank, VEuPathDB, and WormAtlas all began collecting and disseminating data that otherwise would have remained unavailable years before current data sharing requirements. They each use expertise to assess quality, establish and enforce consistency, and apply granular metadata relevant to the community of users. For example, MorphoBank, which publishes data matrices used to compare physical characteristics across species, makes possible the creation of “supermatrices” through the addition of new taxa and features to previously published matrices. This saves researchers from having to recollect samples or manually extract data from articles (O’Leary and Kaufman 2011). WormAtlas provides an extreme example of data rescue where the database contains digital scans of thousands of physical prints and negatives of nematode anatomy created in the 1970s and 80s, when journal page limits meant only a tiny fraction of images made it into the literature. Latent value saw the slides retained, handed between researchers and subsequently moved from lab to lab before being digitized and made available online in the early 2000s (Schroeder and Hall 2021; White 2018). Digitization to improve data access also features in the BHL’s origin story. While centuries of biodiversity materials have been cataloged as part of library or museum holdings, the initiative provided a platform to enable more equitable web access with a corresponding impetus to digitize tens of millions of pages to further enrich the online collection (Freel et al. 2008; *The Biodiversity Heritage Library Project* 2005).

BHL’s example of supercharging the use of existing data collections was found to parallel other databases, including DB1, OMIM, ORDB, rrnDB, and VEuPathDB. Each of these resources provide a focused set of data harvested, in part, from external data resources. The value of centralized access to data relevant to a specific research area emerged as a consistent theme across the documentation for all 9 resources. Databases that make substantial use of external data sources were found to exist in order to lower the barrier to data access—as well as lowering the barrier to data use—by being “fit-for-purpose”, i.e., both reliable *and relevant* to the research question (Gatto et al. 2022). Databases that import and integrate external data work to achieve fit-for-purpose in many ways that are not available from the original data source, for example, through expert annotations (OMIM, VEuPathDB, WormAtlas), authoritative interpretation and summarization (DB1, OMIM, VEuPathDB), and custom analysis tools (BHL, DB1, OMIM, ORDB, rrnDB, and VEuPathDB). Databases containing de novo data, such as MorphoBank and WormAtlas, also provide value-adds that contribute to fit-for-purpose, although they have the added challenge of obtaining the data itself since they are unable to rely on bulk data ingest from existing data sources.

### From Launch to Longevity

The momentum that launched each resource took various shapes but required significant leadership and social capital. In 6 of the cases (DB1, MorphoBank, OMIM, ORDB, rrnDB, WormAtlas), a single champion key to the resource’s inception could be identified. While these individuals often worked in close collaboration with others, they provided (and in a few cases continue to provide) years of leadership directing resource development and operations, while also securing funding. Of these, 4 initial champions have handed over leadership, the most recent being WormAtlas in 2024 while OMIM’s leadership transitioned a few times, with the last occurring in 2002.

BHL, DB2, and VEuPathDB were established under a different dynamic. VEuPathDB, today a suite of 14 databases sharing management and infrastructure, emerged after several individuals launched a pathogenicity-focused effort at NIH’s NIAID. This resulted in the launch of NIAID’s Bioinformatics Resource Centers (BRCs) for Infectious Diseases in 2007, and after a series of mergers, BRCs dedicated to eukaryotic pathogens evolved into VEuPathDB (Amos et al. 2022; Greene et al. 2007). Here elements of individual leadership were clearly critical (such as that of David Roos for one of the original BRCs, ApiDB), but it is less clear that the resource would not have launched at all without any one individual’s participation, i.e., collective interest appears to have been the key source of momentum. DB2’s origin story is similarly rooted in collective interest, as is BHL’s, although the latter came in the form of organizational interest rather than from individual people. Following a series of convening meetings, BHL was established by 10 organizations, including natural history museum libraries, botanical garden libraries, and associated technology partners, who were all interested in making their individual collections more discoverable and usable. These organizations established a consortium that ultimately launched the BHL and then implemented a membership model once initial grant funding ended in 2012 (Freel et al. 2008; Kalfatovic et al. 2023).

Technical transitions were referred to frequently in documentation and interviews despite this topic not being explicitly targeted during the collection of primary source material. The youngest database in this study is 18 years old, long past the typical lifespan of an online biological database (Attwood et al. 2015), so it is not surprising that all 9 resources have undergone major technical updates. Non-exhaustive examples included developing APIs (MorphoBank, OMIM), rewriting the code base (DB2, MorphoBank, rrnDB, VEuPathDB), cloud provisioning (BHL, DB1), and migrating data and servers to new host organizations (MorphoBank, ORDB, rrnDB, WormAtlas, with BHL’s migration forthcoming). While evidence tracing each database’s technical history was not purposefully collected during this study, the readiness of these observations reflect the effort and care required to maintain these resources over the long-term.

Similarly, in depth case studies would be required to trace each database’s detailed funding history, but it’s clear that resources have received monetary support in a variety of ways throughout their existence. For example, some received dedicated funding to launch the resource (e.g., MorphoBank, ORDB), continue development/operations (e.g., OMIM, VEuPathDB), and develop specific features or support initiatives (e.g., BHL, MorphoBank). Other times resource development/operations were one of many specific aims within a research proposal, but not the entirety of it (e.g., WormAtlas). In yet other cases, a database was necessary to address a research question and were conceptually a form of custom-built, in-house equipment that was then made openly available, although not explicitly articulated as an aim or product (e.g., rrnDB). Within the primary source material for each database, especially articles, history pages on current websites, and WayBack Machine snapshots, there was evidence that funding circumstances changed over time such that any one of these scenarios can describe a given resource at various points in its history. These trends are consistent with the experiences of social science data repositories (Eschenfelder et al. 2022).

### Current Funding and Support

For the databases in this study, 7 of the 9 databases currently receive some form of dedicated monetary support, defined as funds explicitly awarded to carry out the work of operating, maintaining, or upgrading the resource. Dedicated monetary support came in several flavors: 1) grants or contracts from US federal agencies or private philanthropies/foundations for a given period of time, 2) membership fees, 3) subscription fees, and 4) donations. Additionally, all 9 databases were found to rely on non-monetary support, which could be broadly categorized as host support and distributed support.

#### Dedicated Monetary Support

As of August 2025, the majority of databases were receiving or are expecting to soon receive dedicated monetary support. This includes 3 databases, MorphoBank, OMIM, and WormAtlas, that were actively receiving dedicated funding from US federal agencies, as well as the 2 anonymous resources that had received federal funding in recent years but elected to not publicly disclose details at the time of writing. Of these, none operate entirely on this funding. Morphobank’s current NSF EAR grant (in a one year no-cost extension) and a recently ended DBI grant were explicitly awarded to upgrade the database architecture and transition to fully self-sustaining at Phoenix Bioinformatics, an independent nonprofit organization established in 2013 to make scientific data widely available and accessible. Another resource, VEuPathDB, was receiving dedicated financial support from NIH NIAID, which covered 60% of its operating costs, but this funding was not renewed in September 2024 (Wadman 2024). VEuPathDB is currently operating on stopgap funding from philanthropic and foundation organizations and additional host support, and they began implementing a new funding model in 2025 after quickly gathering community input. VEuPathDB’s website contains detailed information on their user survey, operating expenses, and the new funding model, which relies on tiered subscriptions based on project budgets and frequency of usage. On its move to Phoenix Bioinformatics, MorphoBank adopted a voluntary membership model in an effort to continue to offer data openly instead of having to restrict access to subscription holders. At the time of writing, database representatives reported still working to collect enough memberships to fully meet their sustainability needs. Memberships are BHL’s main source of dedicated monetary support, a model that’s been in place for over a decade and confers governance rights to member organizations and other benefits that have changed over time, such as the ability to apply for centralized digitization funds. Donations are welcomed on 5 resource websites (BHL, DB1, MorphoBank, OMIM, and VEuPathDB), although no interviewees indicated donations reliably contributed to sustainability, consistent with OECD’s assessment that donations are “a bonus rather than an expectation” (OECD 2017). One interviewee shared that donations bring in “a little money, and it certainly raises community awareness.”

#### Host Support

Hosts provide a wide variety of supports, such as administrative activities associated with fiscal sponsorship, access to IT resources, and the ability to support shared staffing. While these supports have underlying monetary costs, because they are possible through the affordances of scale, which is itself dynamic, they are difficult to efficiently track and accurately quantify per resource. Academic institutions serve as the most common primary host organization (7/9), and each provides both fiscal sponsorship and a technical home for the database’s infrastructure. Phoenix Bioinformatics, MorphoBank’s non-profit host organization, similarly supports both its fiscal and technical operations. As the BHL begins its transition away from the Smithsonian Institution, it recently announced that its new fiscal sponsor is the Council on Library and Information Resources (CLIR), a “independent, nonprofit organization that forges strategies to enhance research, teaching, and learning environments in collaboration with libraries, cultural institutions, and communities of higher learning.” As of writing, efforts to identify support for other activities were underway but had not yet been identified.

Database representatives in this study reported marked variability in the extent of host support available beyond the administrative activities associated with fiscal sponsorship. At a minimum, participants reported support of database leadership through employment of faculty members who devote a fraction of their time towards direction and oversight. At the other end of the spectrum, the Smithsonian Institution’s support of the BHL is substantial, including both technical and administrative staff as well as hosting the technical infrastructure. Other resources reported similar support through departmental or centralized resources that pool technical infrastructure (such as web hosting and storage) and staff (such as shared developers). In some cases (DB2, ORDB, rrnDB, WormAtlas), this support was reported by participants as available through centralized services funded in part through indirect costs, such as IT units and research institutes.

#### Distributed Support

Interviews and documentation provided insight into a variety of forms of community support that were non-monetary and not provided by the host organization, yet are crucial to the database’s operations, especially in aggregate. One example is volunteer service to governance, such as participation on advisory boards and committees (BHL, MorphoBank, VEuPathDB, WormAtlas). Uniquely, ORDB receives periodic volunteer technical support from an early developer of the resource who has since moved on in their professional career but remains interested in the welfare of the database, with the database representative interviewed acknowledging that the volunteer’s priority needs to be “earning a living before we do any of this.” Of the databases within this study, ORDB operates the most leanly, which is possible through tight scoping, early adoption of expert-informed automation, and the personal commitment of a few individuals.

The most common type of distributed support comes via the contribution of data itself. DB1, DB2, MorphoBank, VEuPathDB, and WormAtlas all receive deposits from researchers directly and interviewees noted that they rely on the time and effort it takes to generate data to begin with as well as prepare it for deposit. BHL, DB1, ORDB, OMIM, rrnDB, and VEuPathDB all have automated or semi-automated workflows to ingest relevant data from external sources like the GenBank, the Global Names Architecture, the Kyoto Encyclopedia of Genes and Genomes, Swiss-Prot, and many others—all possible themselves through efforts across the world. BHL aggregates a wide variety of materials, from contemporary born-digital materials to digitized 19th century field notes, through the collection and digitization efforts of its member organizations. Additionally, resources that provide expert interpretation of and reference to peer-reviewed literature, like OMIM, or create comprehensive educational materials, like WormAtlas, depend on the availability of the literature itself. Collectively, these distributed supports depend on the continued health and vitality of life sciences research.

### Impacts

Database representatives were asked about the potential impacts if their data resource had to be shuttered due to a decline in funding, and a wide variety of ramifications were noted – from broken DOIs, to a reduction in their institution’s reputation, to users having to bear the costs of access. There were three strong common themes expressed by participants: higher barriers (9/9), the loss of expertise (7/9), and lost or interrupted opportunities (7/9).

#### Higher Barriers

Many comments coalesced into issues best described as higher barriers for both accessing and using data. These were often discussed in terms of efficiency, concisely summarized in VEuPathDB’s recent User Impact and Sustainability Survey as “having to revert to manual, fragmented, or less accurate methods to gather data, leading to inefficiencies and delays.” Similar anticipated impacts were shared for BHL, DB1, DB2, MorphoBank, OMIM, rrnDB, WormAtlas. The representative for WormAtlas illustrated this by describing how the majority of the *C. elegans* research uses a pipeline that begins with mutagenesis to identify the genes associated with a specific phenotype. Once the gene is found, the presence of the expressed protein must be pinpointed to a specific tissue and even to an individual cell, which researchers do through the precise and detailed anatomical resources provided in WormAtlas. The database representative shared that this mapping is considered so essential that today it’s hard to publish a study without it, and if researchers must perform their own individual anatomical studies in the future, “it’s just going to become less efficient for them.”

Concerns about barriers were often in the context of more detailed and subtle observations that aligned with a concept articulated by former Research Data Alliance secretariat, Lynn Yarmey—the notion of separating the data from the service. This played out in the following way: Several representatives made explicit reference to the impact of data loss should their database go dark. For example, one resource noted that the data within was unique and kept the field grounded in actual data, which was especially valuable for validating (or refuting) claims of novelty or effectiveness (DB1). In another example, the participant expected that an especially useful type of data may become available only through ad hoc networks, both reducing access and appropriate governance (DB2). VEuPathDB’s user survey again echoed these concerns, citing disappearance of specialized data as one of many issues should the resource cease to be available.

However, many representatives also stated that the data itself would continue to exist in some form if the resource went dark, but the barrier to access and use would become prohibitive. WormAtlas, for example, described how their anatomical maps would need to be recreated by combing through hundreds of articles, concluding, “In theory, it’s findable. In principle, it’s hard to do, and it’s not very efficient.” Similarly, BHL shared “The data is backed up, it exists in the IA [Internet Archive]. However, IA, as we’ve seen, is at risk from attack. Also, it is not curated, so finding things in IA is very complicated. It doesn’t mean you can’t. But it is tricky.” MorphoBank’s representative commented “Let’s say the data gets dropped completely, and you can’t even find it in an archive anywhere. People will have to go back to the old way of hunting down supplemental data and transcribing information from old PDFs. Maybe, if you’re lucky, you can find the files.” That is, conceptually, these individuals separated the data from their ultimate concern, which is that expertly gathered data for a known user community is a valuable service in and of itself. Moreover, such collections are not diminished by use–a key characteristic of a public good. This came out even more strongly as others considered impacts if their resource was shuttered. OMIM, which combines several dozen external resources with expert annotation and interpretation, emphasized its role as the starting place for understanding a relationship between genotypes and phenotypes, explaining that there is no comparable resource available. The interviewee shared that “The labs really depend on what’s in OMIM. The research pipelines that try to find new disease genes depend on what’s in OMIM.” This was supported by the website metrics, with tens of thousands of visitors each day (Hamosh et al. 2021), and direct interaction with users, described as “If they [the researchers] read a paper, the first place they go to is OMIM. What did OMIM do? And if we haven’t dealt with that paper yet, they ask—don’t you want to look at this paper? Because they’re waiting to see what we’re going to do.” The service that centralizes and interprets what is otherwise distributed and disparate surfaced as the core concern. VEuPathDB learned this first hand in fall 2024 as cuts began, and they launched their user survey in response. The interviewee described a flood of emails asking what had happened and “How can we survive without it?”. Their survey received 1,862 responses, 80% of which reported weekly usage. VEuPathDB’s representative commented “Science is very resilient. We will recover eventually, but there will be a major impact certainly.” It appears that a large part of this impact is how much researchers rely on the aggregation of relevant data plus associated tools and resources that were developed specifically for the communities to do their work. Without these, participant comments repeatedly emphasized how theoretically available data would go unutilized. There is, in practice, a very real difference between availability and usability.

#### Loss of Subject Matter Expertise

Amongst many of the database representatives, and tied to the loss of service, was a deep concern about the loss of expertise (BHL, DB1, DB2, OMIM, rrnDB, VEuPathDB, WormAtlas). As long-lived resources, many have full- or part- time staff such as database developers, curators, and managers with years, and even decades, of experience. This experience straddled domain and technical knowledge as well as service provision and operations. Interviewees’ concerns reflected plain humanity, given that some had “devoted all their career” to the resource, as well as profound impacts on the resources themselves. In addition to keeping technical infrastructure operational, interviewees reported that the loss of staff expertise would mean the inability to assess quality, uphold standards, interpret data, compare studies, and train new generations of scientists. Moreover, the impact of even a short-term funding crisis made the circumstances especially concerning given that infrastructure itself can withstand a period of low-level maintenance, but experts require paychecks—and steady ones at that. For some, the loss of expertise meant ultimately reverting to a past of inefficiencies, confusion, and disagreements in the field, with one participant sharing that they anticipated deterioration in how the field interprets studies within a few years if their database was shuttered. Interpretation also surfaced in the value of collaboration between experts, with OMIM sharing a typical scenario as “If we’re confused about something, we all come out in the hallway together … everybody go read this paper, let’s sit down, because I think this is a paradigm shift, but I don’t understand how it works. Let’s figure it out.” A team of experts who could freely discuss and debate was considered essential for accurate and nuanced interpretations. VEuPathDB further emphasized the role of developing future expertise. Although the representative expressed long-term confidence in the resilience of science, they saw on the horizon a void in training without VEuPathDB as a centralized source of knowledge to pass on from one scientific generation to the next, essentially resulting in what they described as a “lag phase” in the development of new expertise in pathogenicity. That’s not to say representatives were single-mindedly focused on their resource’s survival; several acknowledged the potential to improve an imperfect system, but they understood and could speak to the direct implications on their user communities with authority and, overall, were especially concerned with cascading ramifications of abrupt loss of expertise.

#### Lost or Interrupted Opportunities

Database interviewees also reported that future visions and plans for responsible and forward-thinking infrastructure management would be thwarted if their resources had to be shuttered or even just left in limbo for an extended period of time. Lost opportunities included a hampered ability to ingest or integrate new data, for example, phenotypes for non-coding RNAs in OMIM, additional species in WormAtlas, or species occurrence data for the BHL. While others were concerned about impacts on planned backend and frontend infrastructure updates without the funding needed to carry out the work. These issues were especially salient given that BHL, DB2, MorphBank, and OMIM all shared that AI bots were aggressively scraping their sites, in some instances taxing the infrastructure to the point where measures had to be taken to deter parasitic practices. Each otherwise was either actively taking steps to incorporate AI into operations themselves or reported interest in partnering with companies for symbiotic AI use of the database’s content. These were also tied to efforts to optimize cost, for example, improving workflows for data deposit or migrating to scalable storage systems. Such efforts also take resources, with MorphoBank sharing that their viability was only possible through NSF grants that supported updating their infrastructure and enabled migration to Phoenix Bioinformatics.

A unique observation on lost opportunities came from rrnDB, which was developed to address a fundamental question – why would any organism have multiple copies of ribosomal RNA? What started with a single investigator’s curiosity evolved into a well-used community resource, with its associated *Nucleic Acids Research* Database Issue papers collectively cited nearly 3000 times to date. This is all the more notable given that data citation is known to dramatically underreport actual usage (Silvello 2018). Robust use comes via those studying microbial communities, like the gut microbiome, because species-level RNA copy number is needed for accurate population sampling. Beyond its concrete, practical utility, the interviewee also noted that the resource “spawns making observations that would be the first step in science,” coming up with an example on the spot of comparing copy number between aquatic versus terrestrial populations. In their view, shuttering the resource meant a lost opportunity to ask questions and interrogate data by reusing infrastructure already in place. While the source data would remain available, rrnDB developed an automated pipeline to (among other things) aggregate by genus, which is the relevant measure for microbial population studies, when genomic sequences are provided at the variant level. The representative noted that the recreation of such a resource from scratch would likely be too labor intensive for individual projects to undertake or, at the very least, substantially slow the progress of their research by having to recreate what rrnDB has already done.

#### Current Impacts

While this study was largely intended to anticipate the potential impacts of new funding challenges, real impacts had already begun by the time interviews were held in summer 2025. Confusion and uncertainty were clearly making it even harder to provide leadership for resources that have long been under pressure. Understanding the risks associated with even temporary funding losses, database representatives reported redirecting time and energy towards investigating alternative funding sources, including non-federal funding sponsors and/or fees for some types of access or support (BHL, DB1, MorphoBank, OMIM, rrnDB, VEuPathDB, WormAtlas). This was also proving more difficult in light of another current reality that became evident—increased competition for funding not only within the grant agencies themselves but other sources. Interviewees and news outlets both report that private funding sources are experiencing extraordinary demand (Glenza 2025; Lenharo 2025). Institutional resources are often supplemental and even more time-limited (OECD 2017). This source may be especially useful for stopgap funding but has also been impacted as organizations brace for challenges across higher education in the US. For example, rrnDB is currently experiencing impacts because they utilize host support for a developer whose salary is partially covered on another grant that was abruptly cancelled, thereby jeopardizing their position and support for rrnDB entirely. A confounding issue is that internal funding, which used to be available in various forms, has grown scarce as “things have tightened up dramatically.”

### Limitations

Because this study aimed to investigate whether long-lived life science databases are particularly vulnerable at this point in time, it was executed on a short time scale by a sole individual. This limited the depth of information gathered for each resource; in-depth, longitudinal case studies would likely reveal fine details and caveats that would provide a more fulsome and nuanced comparison across databases, especially given their variability. As with any interpretive case study, generalizability is limited; here in particular to how observations relate to newer resources, which may have less developed user bases and little institutional memory for weathering hardships. A final limitation is the bias of the author, who is predisposed in favor of data sharing, albeit with constraints. The author heads a unit that facilitates data sharing at a major US research university and, knowing the challenge of sustainability, routinely advises against the launch of brand new open data resources. However, as a trained biochemist, the author remains acutely aware of the crucial role databases play in life-sciences research and is in favor of developing a healthier ecosystem.

## Discussion

Long-lived resources were targeted for this study because any online database requires ongoing care and maintenance, indicating that their continued existence has been purposeful. In fact, the databases in this study have continued to exist not just with purpose, but with persistence in the face of changing leadership, host institutions, technology, and funding. The possibility that resources may “simply disappear” in the wake of new funding cuts was voiced by several data representatives, yet having already weathered challenges in the past, most expected new challenges and were preparing to meet them. With BHL and VEuPathDB already experiencing significant reductions in support, rrnDB the unlucky recipient of cascading effects, MorphoBank still looking to fully meet minimum sustainability goals, a decade-long struggle for funding in the 2010’s still fresh for OMIM, and, external to this study, FlyBase’s recent announcement of cancelled funding, it seems wise for database leadership to ready themselves.

With a past, present, and future seemingly telling a consistent story—funding for databases will be scarce—alternate funding options must be considered in light of past and current challenges. Life science researchers have historically baulked at paying for access to collections of data (National Research Council 2003; Strasser 2019). For example, controversy erupted from the scientific community in 2001 when Celera set up fees to access human genome data. Yet NIH leaders saw a clear and unproblematic parallel to academic publishing, with a report for *Science* in 2001 quoting then director of the National Institute of Mental Health Steven Hyman as saying that he saw “no problem in paying for the data twice: We do it all the time”, further adding, “We pay for the research, we pay for publication costs, and then we pay for the journal subscription for our scientists. We do it without complaining…” (Marshall 2001). However, complaints about the cost and exclusivity of the commercial publishing industry have steadily mounted over the past two decades, with NIH recently requesting comments on plans to limit article publishing charges (NIH, 2025). This suggests that today neither researchers nor the federal government will be complacent paying anything above marginal costs.

This leaves not-for-profit models, with mixed stream revenue being the most likely for success (OECD 2017). Each database covered here already has experience, to greater or lesser extent, diversifying support overall, and most have experience diversifying funding streams. In a future bereft of federal funding, the relevant ratio would need to rebalance significantly. The most natural redirection would be to shift to private philanthropic or foundation support. In the short term, this may be especially difficult as private funders are swarmed with need, but a recent piece by former NSF director and current president of the Science Philanthropy Alliance France Córdova argued one of three key priorities for philanthropic funding is: “Preserving data infrastructure isn’t the flashiest cause, but it’s vital; when data is lost or disrupted, the coherence and reliability of the entire research ecosystem begin to erode.” (Córdova 2025).

Regardless, it’s highly unlikely that philanthropies will be able to make up entirely for the withdrawal of federal funding. Two other common funding streams are referred to as “deposit- side” and “access-side” by OECD. Deposit-side funding, either through individual data publication charges or organizational fees that cover all deposits for an organization, generally requires a strong incentive, or even a mandate, to contribute to a specific resource. Access-side models, also called “user pay” models, are viable when there is both a willingness and capacity to pay at the organization, group, or individual level. While the former condition of willingness has long been questionable, the latter condition of capacity seems poised to be equally uncertain, or more, if the number of new solicitations, awarded grants, stalled grants, and indirect cost recovery remains as unpredictable as seen during 2025. Regardless, for databases dedicated to providing open data for community benefit, the need to charge users is a painful consideration since this directly contributes to inequity and discourages data use, including curiosity-driven, exploratory use. Economists note that this model is generally better aligned with value-add services whose use creates draws on the system and reduces net welfare, such as computationally-intense use, and therefore do not typically qualify as public goods. The computational demands of AI bots seem applicable here. To balance open data principles with economic reality, access-side fees for “rivalrous” services are more often used in concert with some amount of cost-free access to public good data and/or services.

Given current circumstances in the US, it is important to underscore that each of these streams are still vulnerable to wide-spread cuts to research funding and higher education as a whole. This makes the need for diversification more critical than ever if resources are to maintain operations, let alone evolve to continually meet needs. However, shifts in funding require capacity and skill-building to develop detailed and well-justified “business models”, with articulations of “value proposition” and “cost drivers.” In essence, this can appear akin to mission drift as efforts shift from focusing entirely on what researchers need to use the resource effectively to what the database itself needs in order to continue existing for researchers. This is not impossible, but it does require careful balancing.

A last point of discussion is a seeming disconnection in regards to the value of expertise. Each database has something valuable and rare—a focused collection of expertise, applied to data and provided as a service, that have been years in the making. AI is well known for requiring massive amounts of resources, to the alarm of many, but here is a resource that others are fighting to continue to provide in centralized, standardized, and open ways—high-quality, expertly curated data. In fact, at a high level, it seems that AI’s needs and researchers’ are aligned, and there should be a path forward for both needs to merge in a balanced and truly synergistic way. The contours of such partnerships would require tough negotiation.

## Conclusions

For long-lived databases, the current funding climate is not unfamiliar, but it is more pressing as resources previously considered supplemental are looked to for primary sources of funding. The dissonance is also not especially new. Data sharing and reuse have been nation-wide priorities at a policy level for over a decade in the US, but funding for data collections continues to be hard-fought. An ever-growing number of online resources, should they all seek to live on in perpetuity, was never sustainable, and the current funding cuts in the US make the need for cohesive strategies and a long-term vision more important than ever.

The open data mantra “as open as possible, as closed as necessary” was meant to account for privacy, security, and intellectual property considerations (Landi et al. 2020). However, for more and more databases, a more realistic and accurate mantra may be “as open as possible, as closed as self-sustaining requires.” Database leaders exhibit a public service mindset, and as one interview wrapped up, the conversation morphed into succession planning. The participant observed that they have had “an inherent motivation to do it, but if I ask somebody else, they might say ‘all I see is headaches’. I could say, ‘but they’re small headaches’, but then I don’t see many advantages [for someone to take on the responsibility].” While incentives have existed to remain operational to date, the incentives may evaporate as headaches continually mount and leaders are pressed into adopting funding models that clash with sudden change, economic realities, fundamental principles—or all three.

## Supporting information

Table S5. Database Profiles

Code definitions and examples

## Acknowledgements

The author wishes to thank Kelli Trei (BHL), Brooke Long-Fox and Tanya Berardini (MorphoBank), Ada Hamosh (OMIM), Chiquito Crasto (ORDB), Tom Schmidt (rrnDB), George Christophides (VEuPathDB), Nate Schroeder (WormAtlas), and two anonymous participants for their time and insights. The author also wishes to thank the generous donor support as the Allen and Elaine Avner Professor in Interdisciplinary Research and Illinois Computes for MAXQDA licensing, as well as Sandi Caldrone and MH Han for providing feedback on the manuscript.

## Conflict of Interest

During the preparation of the final manuscript, the author accepted an invitation to serve on the WormAtlas advisory board and was previously consulted during migration of the resource to the University of Illinois Urbana-Champaign. The author also previously consulted with the Global Biodata Coalition on a project to create a large-scale inventory of biodata resources.

