## Supplemental Figure S1 and Tables S1-4 for "Sustainability During Instability: Long-Lived Life Science Databases and Science Funding Outlook in the United States"

### Supplemental Materials

#### Figure S1. Code Matrix.

Code Matrix exported from MAXQDA 24 for all codes used in the study. Unweighted analytical codes with hits counted once per document, colored by group, and circle size calculated per row. Total sums for code occurrence included per code (last column) and per unit (last row).

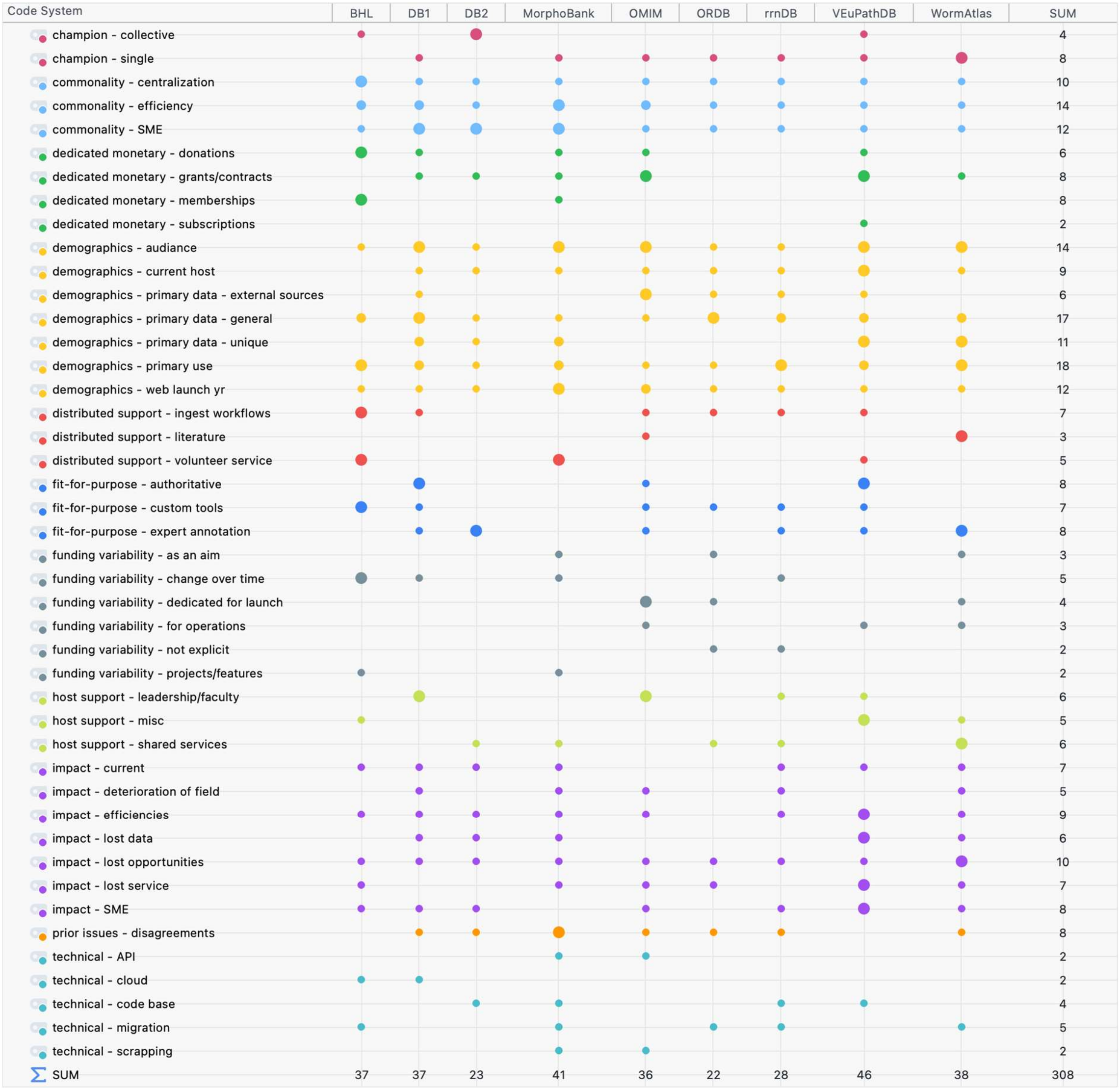

**Table S1. Primary Source Material Counts per Database**

| Database | Interviews | Webpages | Articles | Other Documents (e.g., reports, news media, etc.) |
|---|---|---|---|---|
| BHL | 1 | 7 | 3 | 1 |
| DB1 | 1 | 3 | 1 | 3 |
| DB2 | 1 | 4 | 1 | 0 |
| MorphoBank | 1 | 5 | 2 | 4 |
| OMIM | 1 | 3 | 3 | 0 |
| ORDB | 1 | 2 | 2 | 0 |
| rrnDB | 1 | 3 | 3 | 0 |
| VEuPathDB | 1 | 9 | 8 | 2 |
| WormAtlas | 1 | 5 | 2 | 1 |
| **Total** | **9** | **41** | **25** | **11** |

##### Table S2. Primary Source Materials

| Unit | Document Name | Word Count | Access* |
|---|---|---|---|
| BHL | Webpage - BHL Training | 351 | http://web.archive.org/web/20250828235403/https://about.biodiversitylibrary.org/ufaqs/where-can-i-find-training-materials-about-using-bhl-collections-and-services/ |
| BHL | Webpage - BHL Tool | 878 | http://web.archive.org/web/20250514105002/https://about.biodiversitylibrary.org/ufaqs/how-does-the-taxonomic-name-recognition-algorithm-work-in-bhl/ |
| BHL | Webpage - BHL Memberships as of 2012 | 1160 | http://web.archive.org/web/20250614145150/https://blog.biodiversitylibrary.org/2016/05/biodiversity-heritage-library-adds-bhl-australia-as-a-new-member.html |
| BHL | Webpage - BHL History | 1266 | http://web.archive.org/web/20250704150445/https://about.biodiversitylibrary.org/about/history-of-bhl/ |
| BHL | Webpage - BHL Funding | 717 | http://web.archive.org/web/20250814210255/https://about.biodiversitylibrary.org/about/bhl-funding/ |
| BHL | Webpage - BHL Field Notes | 9528 | http://web.archive.org/web/20241204065810/https://www.biodiversitylibrary.org/collection/FieldNotesProject |
| BHL | Webpage - BHL About | 726 | http://web.archive.org/web/20241204184328/https://about.biodiversitylibrary.org/ |
| BHL | Video - BHL | 8047 | Not Available |
| BHL | Document - BHL prospectus 2005 | 1394 | https://web.archive.org/web/20250328083718/https://www.sil.si.edu/bhl/supportdocuments/BHLP-prospectus10-05.pdf |
| BHL | Article - BLH Code4Lib 2008 | 3335 | https://web.archive.org/web/20250329102814/https://journal.code4lib.org/articles/52 |
| BHL | Article - BHL Tools 2019 (Abstract) | 880 | https://doi.org/10.3897/biss.3.35353 |
| BHL | Article - BHL BISS 2023 (Abstract) | 896 | https://doi.org/10.3897/biss.7.112430 |
| DB1 | Webpage - DB1 3 | redacted | Not Available |
| DB1 | Webpage - DB1 2 | redacted | Not Available |
| DB1 | Webpage - DB1 1 | redacted | Not Available |
| DB1 | Interview Notes - DB1 | redacted | Not Available |
| DB1 | Document - DB1 3 | redacted | Not Available |

| | | | |
|---|---|---|---|
| DB1 | Document - DB1 2 | redacted | Not Available |
| DB1 | Document - DB1 1 | redacted | Not Available |
| DB1 | Article - DB1 | redacted | Not Available |
| DB2 | Webpage - DB2 4 | redacted | Not Available |
| DB2 | Webpage - DB2 3 | redacted | Not Available |
| DB2 | Webpage - DB2 2 | redacted | Not Available |
| DB2 | Webpage - DB2 1 | redacted | Not Available |
| DB2 | Interview Notes - DB2 | redacted | Not Available |
| DB2 | Article - DB2 | redacted | Not Available |
| Morphobank | Webpage - MorphoBank WBM 2007 | 160 | https://web.archive.org/web/20091004075255/http://www.morphobank.org/index.php?g=about&s=more |
| Morphobank | Webpage - Morphobank Home | 258 | https://web.archive.org/web/20250501011441/https://morphobank.org/ |
| Morphobank | Webpage - MorphoBank History | 970 | https://web.archive.org/web/20250927231049/https://phoenixbioinformatics.atlassian.net/wiki/spaces/MD/pages/42230791/About+MorphoBank |
| Morphobank | Webpage - MorphoBank FAQ | 8270 | https://web.archive.org/web/20250927231220/https://phoenixbioinformatics.atlassian.net/wiki/spaces/MD/pages/42226561/FAQ |
| Morphobank | Webpage - MorphoBank Education | 9570 | http://web.archive.org/web/20250209075843/https://phoenixbioinformatics.atlassian.net/wiki/spaces/MD/pages/42226624/User+Guide+-+Overview |
| Morphobank | Video - MorphoBank | 7297 | Not Available |
| Morphobank | Document - MorphoBank Workshop 2001 | 3175 | https://web.archive.org/web/20071006171557/http://morphobank.geongrid.org/articles/morphobank_workshop_report.pdf |
| Morphobank | Document - MorphoBank Supp Comments | 26 | Not Available |
| Morphobank | Document - MorphoBank OLeary CV | 6836 | https://web.archive.org/web/20240717183004/https://renaissance.stonybrookmedicine.edu/system/files/O%27Leary%20full%20Cv.pdf |

| | | | |
|---|---|---|---|
| Morphobank | Document - MorphoBank NSF 9903964 | 910 | https://web.archive.org/web/20250224005603/https://www.nsf.gov/awardsearch/showAward?AWD_ID=9903964&HistoricalAwards=false |
| Morphobank | Article - MorphoBank Planning 2003 | 4708 | https://doi.org/10.1126/science.300.5626.1692 |
| Morphobank | Article - MorphoBank Cladistics 2011 | 6791 | https://doi.org/10.1111/j.1096-0031.2011.00355.x |
| OMIM | Webpage - OMIM FAQ | 2590 | https://web.archive.org/web/20250604110638/https://omim.org/help/faq |
| OMIM | Webpage - OMIM External Links | 1120 | https://web.archive.org/web/20250813005320/https://www.omim.org/help/external |
| OMIM | Webpage - OMIM About | 296 | https://web.archive.org/web/20250507082318/https://www.omim.org/help/about |
| OMIM | Video - OMIM | 13479 | Not Available |
| OMIM | Article - OMIM NAR 1994 | 2992 | https://doi.org/10.1093/nar/22.17.3470 |
| OMIM | Article - OMIM BJHS 2021 | 12429 | https://doi.org/10.1017/S0007087421000224 |
| OMIM | Article - OMIM AJMG 2021 | 4644 | https://doi.org/10.1002/ajmg.a.62407 |
| ORDB | Webpage - ORDB History | 922 | https://web.archive.org/web/20240817104542/https://ordb.biotech.ttu.edu/OrDB/info/ordb_versions |
| ORDB | Webpage - ORDB FAQ | 678 | https://web.archive.org/web/20240817094337/https://ordb.biotech.ttu.edu/OrDB/info/ordb_faqs |
| ORDB | Video - ORDB | 11307 | Not Available |
| ORDB | Article - ORDB NAR 2002 | 3962 | https://doi.org/10.1093/nar/30.1.354 |
| ORDB | Article - ORDB Chem Senses 1997 | 3085 | https://doi.org/10.1093/chemse/22.3.321 |
| rrnDB | Webpage - rrnDB Manual | 1659 | https://web.archive.org/web/20250322203000/https://rrndb.umms.med.umich.edu/help/ |
| rrnDB | Webpage - rrnDB Education | 889 | https://web.archive.org/web/20220815152922/https://forum.qiime2.org/t/normalize-asv-by-16s-copy-number/12624 |
| rrnDB | Webpage - rrnDB About | 1278 | https://web.archive.org/web/20250619062152/https://rrndb.umms.med.umich.edu/about/ |

| | | | |
|---|---|---|---|
| rrnDB | Video - rrnDB | 8276 | Not Available |
| rrnDB | Article - rrnDB NAR 2015 | 4779 | https://doi.org/10.1093/nar/gku1201 |
| rrnDB | Article - rrnDB NAR 2008 | 3803 | https://doi.org/10.1093/nar/gkn689 |
| rrnDB | Article - rrnDB NAR 2001 | 2642 | https://doi.org/10.1093/nar/29.1.181 |
| VEuPathDB | Webpage - VEuPathDB Survey Results | 344 | https://web.archive.org/web/20250307164213/https://static-content.veupathdb.org/documents/Survey_Summary.pdf |
| VEuPathDB | Webpage - VEuPathDB Subscriptions | 596 | https://veupathdb.org/veupathdb/app/static-content/subscriptions.html |
| VEuPathDB | Webpage - VEuPathDB Subscribers June 2025 | 2043 | https://veupathdb.org/veupathdb/static-content/subscribers.html |
| VEuPathDB | Webpage - VEuPathDB People | 830 | https://veupathdb.org/veupathdb/app/static-content/personnel.html |
| VEuPathDB | Webpage - VEuPathDB General | 762 | https://veupathdb.org/veupathdb/app/static-content/about.html |
| VEuPathDB | Webpage - VEuPathDB FAQ | 5126 | https://veupathdb.org/veupathdb/app/static-content/faq.html |
| VEuPathDB | Webpage - VEuPathDB Deposit 2010 | 1681 | https://veupathdb.org/veupathdb/app/static-content/dataSubmissionReleasePolicy.html |
| VEuPathDB | Webpage - VEuPathDB Comm | 671 | https://veupathdb.org/veupathdb/app/static-content/acks.html |
| VEuPathDB | Webpage - VEuPathDB About | 1388 | https://veupathdb.org/veupathdb/app/static-content/about.html |
| VEuPathDB | Video - VEuPathDB | 7564 | Not Available |
| VEuPathDB | Document - VEuPathDB Full Survey | 5866 | https://web.archive.org/web/20250404121855/https://static-content.veupathdb.org/documents/PUBLIC_REPORT_VEuPathDB_User_Impact_Sustainability_Survey.pdf |
| VEuPathDB | Document - VEuPathDB Education | image | https://veupathdb.org/veupathdb/app/static-content/landing.html |
| VEuPathDB | Article - VEuPathDB Science 2024 | 2015 | https://doi.org/10.1126/science.zbgrc0p |
| VEuPathDB | Article - VEuPathDB Roos 2019 | 2495 | https://web.archive.org/web/20250514121024/https://www.the-scientist.com/parasitologist-reprogrammed-a-profile-of-david-roos-30015 |

| | | | |
|---|---|---|---|
| VEuPathDB | Article - VEuPathDB Plasmodb NAR 2001 | 3037 | https://doi.org/10.1093/nar/29.1.66 |
| VEuPathDB | Article - VEuPathDB Para Today 1998 | 3057 | https://doi.org/10.1016/s0169-4758(98)01300-3 |
| VEuPathDB | Article - VEuPathDB NAR 2023 | 6748 | https://doi.org/10.1093/nar/gkad1003 |
| VEuPathDB | Article - VEuPathDB NAR 2021 | 10163 | https://doi.org/10.1093/nar/gkab929 |
| VEuPathDB | Article - VEuPathDB NAR 2010 | 2660 | https://doi.org/10.1093/nar/gkp941 |
| VEuPathDB | Article - VEuPathDB Infect Immun 2007 | 7551 | https://doi.org/10.1128/IAI.00105-07 |
| WormAtlas | Webpage - WormAtlas About | 708 | https://web.archive.org/web/20250705233055/https://wormatlas.org/about.htm |
| WormAtlas | Webpage - WormAtlas WBM 2002 | 327 | https://web.archive.org/web/20020405080718/http://www.wormatlas.org/acknowledgements.htm |
| WormAtlas | Webpage - WormAtlas Handbook Hermaphrodite | 409 | https://doi.org/10.3908/WORMATLAS.1.1 |
| WormAtlas | Webpage - WormAtlas Best of The Web 2011 | 618 | https://web.archive.org/web/20240221205846/https://www.genengnews.com/best-of-the-web/worm-atlas/ |
| WormAtlas | Webpage - WormAtlas ACES Annouc 2023 | 604 | https://web.archive.org/web/20240519093924/https://neuroscience.illinois.edu/news/2023-03-17t181559/wormatlas-expanding-beyond-c-elegans-support-nih |
| WormAtlas | Video - WormAtlas | 8198 | Not Available |
| WormAtlas | Document - WormAtlas Supp Comments | 197 | Not Available |
| WormAtlas | Article - WormAtlas White MRC 2018 | 7309 | https://www.ncbi.nlm.nih.gov/books/NBK153591/ |
| WormAtlas | Article - WormAtlas JNem 2021 | 1209 | https://doi.org/10.21307/jofnem-2021-090 |

* DOIs and WayBack Machine links provided when possible

**Table S3. Interview Prompts**

| | Question |
|---|---|
| 1 | *What purpose and communities has [name of participant's database] served?* |
| 2 | *Can you outline how [name of participant's database] was established and what effort it took to establish it as a resource?* |
| 3 | *How is [name of participant's database] currently supported (direct and indirectly)?* |
| 4 | *What impacts do you anticipate from a decline of federal funding to support research?* |
| 5 | *If [name of participant's database] must be shuttered due to a decline in support, what impacts do you expect on the future of research in this area?* |
| 6 | *Are there additional impacts on people or your institution?* |
| 7 | *Do you have additional comments that you would like to share?* |

**Table S4. US Federal Agency Abbreviations**

| Acronym | Full Name |
|---|---|
| NSF | National Science Foundation |
| BIO | Directorate for Biological Sciences (NSF) |
| IBN | Integrative Biology and Neuroscience (NSF) |
| DBI | Division of Biological Infrastructure (NSF) |
| EAR | Division of Earth Sciences (NSF) |
| SABI | Sustained Availability of Biological Infrastructure (NSF) |
| NIH | National Institutes of Health |
| OD | Office of the Director (NIH) |
| NCRR | National Center for Research Resources (closed 2011) |
| NHGRI | National Human Genome Research Institute (NIH) |
| NIAID | National Institute of Allergy and Infectious Diseases (NIH) |
| NIDCD | National Institute of Deafness and Other Communication Disorders (NIH) |
| NIMH | National Institute of Mental Health (NIH) |
| NLM | National Library of Medicine (NIH) |
| IAIMS | Integrated Academic Information Management System (NLM) |
| NASA | National Aeronautics and Space Administration |
