## Supplementary material for "Sustainability During Instability: Long-Lived Life Science Databases and Science Funding Outlook in the United States": Table S5. Database Profiles

<table>
<tr><th rowspan="2">Resource Name<br>▪ URL<br>▪ 1st Yr on Web<br>▪ Current Primary Host</th><th rowspan="2" colspan="2">Foundation for Launch<br>Need – Champion – Funding</th><th colspan="5">Current State – as of August 2025</th></tr>
<tr><th>Data & Sources</th><th>Primary Audiences & Uses</th><th>Dedicated Monetary Funding</th><th>Other Supporting Activities</th><th>Operational Status</th></tr>
<tr><td rowspan="3">BHL: Biodiversity Heritage Library<br>▪ biodiversitylibrary.org<br>▪ 2007<br>▪ In transition from the Smithsonian Institution</td><td>Need</td><td>Access to and use of biodiversity data within literature and archives otherwise distributed across multiple collections</td><td rowspan="3">Data and metadata from high resolution archival (scans) and contemporary (born digital) literature and archival material of biodiversity relevance;<br><br>Sourced from distinct libraries and natural history museums, internal SME</td><td rowspan="3">Biologists, conservationists, taxonomists, educators, artists<br><br>Used as primary source material to study historical trends in biodiversity, verification of taxonomic names and classifications; used for educational and artistic purposes</td><td rowspan="3">• Membership fees<br>• Donations welcomed</td><td rowspan="3">Host Support – Leadership, shared staffing, technical infrastructure, and operations management<br><br>Distributed Support – Volunteer effort for selection, scanning, deposition of literature, participation in governance, funding writing, etc.</td><td rowspan="3">Undergoing major leadership and infrastructure transition due to loss of substantial support from the Smithsonian Institution as of Jan 2026;<br><br>Seeking new sponsorship, actively pursuing other funding streams, may need to consider changes to fees in the future</td></tr>
<tr><td>Champion</td><td>Collective interest from 10 founding organizations following a special convening meeting</td></tr>
<tr><td>Initial Funding & Support</td><td>Public and private funding; hosted at the Smithsonian Institution and Missouri Botanical Garden</td></tr>
<tr><td rowspan="3">MorphoBank<br>▪ morphobank.org<br>▪ 2004<br>▪ Phoenix Bioinformatics</td><td>Need</td><td>Access to and use of morphological data otherwise unavailable or distributed throughout the literature</td><td rowspan="3">Phylogenetic matrices and associated data (taxa, characters and states, cells with or without images, bibliographic citations);<br><br>Sourced from individual deposits associated with peer reviewed publications and internal SME</td><td rowspan="3">Evolutionary biologists, systematists, paleontologists, taxonomists and other biologists<br><br>Used to build on previous research and support large-scale comparative studies</td><td rowspan="3">• Membership fees<br>• NSF EAR (in no cost extension)<br>• Donations welcomed</td><td rowspan="3">Host Support – Leadership, staffing (e.g., expert curators), technical infrastructure, and operations management<br><br>Distributed Support – Data creation and deposition from researchers</td><td rowspan="3">Completed major leadership and infrastructure transition from Stony Brook University to Phoenix Bioinformatics in 2021, made possible by NSF DBI SABI funding;<br><br>Seeking growth in memberships to fully cover operating costs</td></tr>
<tr><td>Champion</td><td>Individual champion followed by community interest</td></tr>
<tr><td>Initial Funding & Support</td><td>Dedicated NSF funding for resource; hosted at Stony Brook University</td></tr>
</table>

**Table S5: Database Profiles, continued.**

| **Resource Name**<br>▪ URL<br>▪ 1st Yr on Web<br>▪ Current Primary Host | **Foundation for Launch**<br>Need – Champion – Funding | | **Current State – as of August 2025** | | | | |
|---|---|---|---|---|---|---|---|
| | | | **Data & Sources** | **Primary Audiences & Uses** | **Dedicated Monetary Funding** | **Other Supporting Activities** | **Operational Status** |
| **OMIM: Online Mendelian Inheritance in Man**<br>▪ omim.org<br>▪ 1995<br>▪ Johns Hopkins University | **Need** | Access to and use of comprehensive knowledge on genes and genetic disorders otherwise distributed throughout the literature and across external data sources | Gene and phenotype entries with relationship mapping, clinical summaries, and references to corresponding literature and external data sources;<br><br>Sourced from the biomedical literature, external data sources, and internal SME | Biomedical researchers, e.g., physicians, clinicians, geneticists, as well as educators and students<br><br>Used as the "first stop" authoritative resource for exploration of genes and genetic disorders;<br>Used in medical education at multiple levels | • NIH NHGRI U41<br>• Donations welcomed | *Host Support* – Leadership<br><br>*Distributed Support* – Data and services from external data and literature resources | Dependent on federal funding;<br><br>Aware that alternate funding streams will be insufficient to sustain the resource |
| | **Champion** | Individual champion followed by US federal agency interest in online access | | | | | |
| | **Initial Funding & Support** | Dedicated NIH NLM funding for online version of catalogue of Human Genes and Genetic Disorders; hosted at Johns Hopkins University | | | | | |
| **ORDB: Olfactory Receptor Database**<br>▪ ordb.biotech.ttu.edu/ORDB<br>▪ 1994<br>▪ Texas Tech University | **Need** | Access to and use of chemoreceptor data otherwise distributed across external data sources and within individual research groups | Nucleotide and amino acid sequences for olfactory receptor (OR) and olfactory receptor-like (ORL) gene and protein sequences with interoperability between research labs and partner databases;<br><br>Sourced from external data sources and individual labs and internal SME | Neuroscientists, computational biologists, molecular biologists, bioinformaticians<br><br>Used to study OR and ORL genes, access to keys for variable lab IDs, and access tools to analyze unpublished sequences | None | *Host Support* – Leadership, student training opportunities, technical infrastructure, and operations management<br><br>*Distributed Support* – Volunteer effort for technical development and data and services from external data sources | Dependent on host and volunteer support for light steady-state maintenance with periodic student-lead improvements;<br><br>No new impacts expected while continually considering new funding opportunities |
| | **Champion** | Individual champion followed by collective interest at society conference meeting | | | | | |
| | **Initial Funding & Support** | Dedicated NIH NLM IAIMS funding for resource, additional from NIH NIMH, NIH NIDCD and NASA; hosted at Yale University | | | | | |

**Table S5: Database Profiles, continued.**

| **Resource Name**<br>▪ URL<br>▪ 1st Yr on Web<br>▪ Current Primary Host | **Foundation for Launch**<br>Need – Champion – Funding | **Current State – as of August 2025**<br>**Data & Sources** | **Primary Audiences & Uses** | **Dedicated Monetary Funding** | **Other Supporting Activities** | **Operational Status** |
|---|---|---|---|---|---|---|
| **rrnDB: ribosomal RNA operon copy number database**<br>▪ [rrndb.umms.med.umich.edu](rrndb.umms.med.umich.edu)<br>▪ 2001<br>▪ University of Michigan | **Need**: Access to and use of data to track and compare microbial ribosomal RNA data to copy number otherwise distributed across external data sources<br>**Champion**: Individual champion followed by community uptake<br>**Initial Funding & Support**: NSF BIO IBN and Michigan State University; hosted at Michigan State University | 16S and 23S RNA genes (sequencing and earlier experimental), organism taxonomies, statistical summaries, additional metadata and links to related external resources;<br><br>Sourced from external data resources and internal SME | Microbiologists, ecologists, researchers studying microbial populations (e.g., microbiome)<br><br>Used for basic research (initial) with subsequent practical use to counteract bias while measuring composition of microbial populations | None | *Host Support* – Leadership, shared staffing, technical infrastructure, and operations management<br><br>*Distributed Support* – Data and services from other distributed data and literature resources | Dependent on host support and facing implications from changes to indirect costs and sharing staffing;<br><br>Seeking additional stopgap host support for shared technical staff whose position is in jeopardy after funding for other half of salary line was cancelled elsewhere in unit |
| **VEuPathDB: Eukaryotic Pathogen, Vector, and Host Informatics Resources**<br>▪ [veupathdb.org](veupathdb.org)<br>▪ 2004<br>▪ Universities of Pennsylvania and Georgia | **Need**: Access to and use of genomic data integrated with extensive biological data of pathogenic relevance otherwise distributed across external data sources<br>**Champion**: Individual champions followed by US federal agency interest in pathogenicity<br>**Initial Funding & Support**: NIH NIAID Bioinformatics Resource Centers for Biodefense and Emerging/Re-Emerging Infectious Disease (BRCs); hosted at multiple locations | Genome sequence and annotation, transcriptomics, proteomics, epigenomics, metabolomics, population resequencing, clinical data, surveillance data, host-pathogen interactions, and orthology profiles;<br><br>Sourced from external data sources, individual deposits, and internal SME | Infectious disease researchers broadly, including parasitologists, vector biologists, epidemiologists, field biologists and clinicians;<br><br>Used for centralized access to multi-omic data and associated specialized tools to explore, analyze, and compare genomic and functional data on pathogens, vectors, hosts, and disease | • Stopgap funding from private foundations and universities<br>• Membership fees just beginning<br>• Donations welcomed | *Host Support* – Leadership, shared staffing, technical infrastructure, and operations management<br><br>*Distributed Support* – Data and services from other distributed data and literature resources | Undergoing major transition due to recent loss of substantial support from NIH NIAID as of Sept 2024;<br><br>Stopgap funding and support secured for short term, has begun to restrict some portions of the resource in order to support a subscription model |

**Table S5: Database Profiles, continued.**

<table>
<tr><th rowspan="2">Resource Name<br>▪ URL<br>▪ 1st Yr on Web<br>▪ Current Primary Host</th><th rowspan="2" colspan="2">Foundation for Launch<br>Need – Champion – Funding</th><th colspan="5">Current State – as of August 2025</th></tr>
<tr><th>Data & Sources</th><th>Primary Audiences & Uses</th><th>Dedicated Monetary Funding</th><th>Other Supporting Activities</th><th>Operational Status</th></tr>
<tr><td rowspan="3">WormAtlas<br>▪ wormatlas.org<br>▪ 2002<br>▪ University of Illinois Urbana-Champaign</td><td>Need</td><td>Access to and use of otherwise unavailable C. elegans microscopy data from key developmental biology research and data distributed amongst individual research groups</td><td rowspan="3">Expertly annotated electron microscopy images organized by species, tissue, genotype, age, or sex with corresponding detailed anatomical maps, handbooks, and interactive educational resources<br><br>Sourced from labs, individuals, and internal SME</td><td rowspan="3">Cell biologists, developmental biologists, geneticists, neuroscientists<br><br>Used to compare and study anatomical features, cellular structures, identification of cellular location of gene expression, developmental processes, and phenotype variations;<br>Used for educational materials and in teaching</td><td rowspan="3">NIH OD Resource-Related Research Projects (R24)</td><td rowspan="3">Host Support – Leadership, shared staffing, technical infrastructure, and operations management<br><br>Distributed Support – Data creation and deposition from researchers</td><td rowspan="3">Dependent on federal funding;<br><br>Completed major transition of infrastructure and leadership from Albert Einstein College of Medicine to University of Illinois Urbana-Champaign in 2024;<br><br>Aware of potential impacts of reduction in indirect funds to support shared resources as well as uncertainty of future grant success</td></tr>
<tr><td>Champion</td><td>Individual champion followed by community uptake</td></tr>
<tr><td>Initial Funding & Support</td><td>NIH NCRR; hosted at Albert Einstein College of Medicine</td></tr>
<tr><td rowspan="3">DB1: Database 1<br>▪ > 15 yrs ago<br>▪ US Organization</td><td>Need</td><td>Access to and use of data otherwise unavailable, distributed throughout the literature, and across external data sources</td><td rowspan="3">Data of biomedical relevance;<br><br>Sourced from public databases, individual deposits, and internal SME</td><td rowspan="3">Biomedical researchers and human health experts<br><br>Used to access high-quality data and tools relevant to the study of human health</td><td rowspan="3">Not disclosed</td><td rowspan="3">Host Support – Leadership<br><br>Distributed Support – Data and services from external data and literature resources, data creation and deposition from researchers</td><td rowspan="3">Dependent on federal funding;<br><br>Aware that commercial funding would not be viable and has begun investigating private and local stopgap funding</td></tr>
<tr><td>Champion</td><td>Individual champion followed by community uptake</td></tr>
<tr><td>Initial Funding & Support</td><td>US Federal Funding Agency; hosted by a US Organization</td></tr>
</table>

**Table S5: Database Profiles, continued.**

<table>
<tr><th rowspan="2">Resource Name<br>▪ URL<br>▪ 1st Yr on Web<br>▪ Current Primary Host</th><th rowspan="2" colspan="2">Foundation for Launch<br>Need – Champion – Funding</th><th colspan="5">Current State – as of August 2025</th></tr>
<tr><th>Data<br>& Sources</th><th>Primary Audiences<br>& Uses</th><th>Dedicated Monetary Funding</th><th>Other Supporting Activities</th><th>Operational Status</th></tr>
<tr><td rowspan="3">DB2: Database 2<br>▪ > 15 yrs ago<br>▪ US Organization</td><td>Need</td><td>Access to and use of data otherwise unavailable or distributed throughout the literature</td><td rowspan="3">Data of biomedical relevance;<br><br>Sourced from individual deposits and internal SME</td><td rowspan="3">Biomedical researchers and human health experts<br><br>Used to access high-quality data and tools relevant to the study of human health</td><td rowspan="3">Not disclosed</td><td rowspan="3">Host Support – leadership, shared staffing, technical infrastructure, and operations management<br><br>Distributed Support – Data creation and deposition from researchers</td><td rowspan="3">Dependent on federal funding;<br><br>Concerned with a wide range of operational challenges, aware that resource’s future is uncertain, has considered sunsetting/transfer as well as models to become less dependent on federal funds</td></tr>
<tr><td>Champion</td><td>Collective interest followed by community uptake</td></tr>
<tr><td>Initial Funding & Support</td><td>US Federal Funding Agency; hosted by a US Organization</td></tr>
</table>
