## Supplementary material for "Sustainability During Instability: Long-Lived Life Science Databases and Science Funding Outlook in the United States": Code definitions and examples

**champion - collective**

initiative of multiple people or organizations launched resource

example: "At the time of its formation, BHL included ten member organizations in the U.S. and the U.K." (Webpage - BHL History, p. 1)

**champion - single**

initiative of a single person launched resource

example: "However, David Hall at the Albert Einstein College of Medicine (Bronx, NY) kept the flame of C. elegans neuroanatomy burning, and obtained a grant to set up a central repository of C. elegans anatomy." (Article - WormAtlas White MRC 2018, p. 13)

**commonality - centralization**

value of having relevant data and tools in a single resource

example: "It is a central, shared resource, like the other international sequence databases. However, it also has features specifically designed for the needs of the olfactory research community." (Article - ORDB Chem Senses 1997, p. 2)

**commonality - efficiency**

purpose of the resource is to make research easier, faster, and/or more thorough

example: "Building supermatrices (de Queiroz and Gatesy, 2007) for robust tests of character congruence often requires adding new taxa and characters to published work. It is inefficient to recollect (or even retype) published phenomic matrix data from scratch, just as it is inefficient to re-sequence a gene previously used for molecular phylogenetics, unless there is reason to suspect error or a need to increase sampling." (Article - MorphoBank Cladistics 2011, p. 530)

**commonality - SME**

role of deep professional knowledge of data, topic, and/or infrastructure built up over time

example: "...that have been vetted and that have been, in many cases, with great effort brought up to standard of containing all the appropriate information." (Video - MorphoBank: 187-188)

**dedicated monetary - donations**

solicitation of spontaneous financial contributions of any amount to support the work of the resource

example: "Can VEuPathDB accept additional contributions, and are such contributions tax-deductible? Yes. Please see Charitable Contributions in support of VEuPathDB." (Webpage - VEuPathDB FAQ, p. 7)

**dedicated monetary - grants/contracts**

funds explicitly awarded to carry out the work of operating, maintaining, or upgrading the resource for a given period of time, either from a government or private agency example: "... we were able to secure a NSF sustainability grant to help us make that transition from pure grant funding to something else." (Video - MorphoBank: 94-96)

**dedicated monetary - memberships**

funds paid, generally on a reoccurring basis, on behalf of an organization to support a resource that is important to the organization's mission or community

example: "When initial grant funding ended in 2012, BHL established an annual dues model for its Members and Affiliates to help support central BHL operating expenses and technical development." (Article - BHL BISS 2023, p. 2)

**dedicated monetary - subscriptions**

funds paid, generally on a reoccurring basis, on behalf a user, group, or organization in order to obtain access to all or part of a data resource

example: "The changing funding landscape makes it difficult to sustain essential infrastructure through grants, necessitating a mandatory subscription service." (Webpage - VEuPathDB Subscriptions, p. 1)

**demographics - audience**

users of the data resource

example: "These users range from high school, college, graduate, genetic counseling, and medical students to residents, geneticists, clinicians in other disciplines, as well as basic, translational, and computational scientists." (Article - OMIM AJMG 2021, p. 4)

**demographics - current host**

organization currently providing fiscal, operational, and/or technical hosting, generally only one

example: "currently running under the auspices of Texas Tech University" (Video - ORDB: 247-248)

**demographics - primary data - external sources**

data ingested from another data resource, such as GenBank, KEGG, etc.

example: "Major Data Sources
Starting with v5.0 the data sources for rrnDB are as follows.
- NCBI genome assemblies and annotation data from the NCBI FTP server.
- NCBI taxonomy data are acquired from the NCBI FTP server.
- RDP taxonomy data are acquired from the RDP Classifier tool of the Ribosome Database Project (RDP).
- Records inherited from rrnDB v3.1.227 are based on empirical determination of rrn copy number using various methods not involving finished genome sequences." (Webpage - rrnDB About, p. 2)

**demographics - primary data - general**

primary data that the resource manages and disseminates

example: "Data types include genome sequence and annotation, transcriptomics, proteomics, epigenomics, metabolomics, population resequencing, clinical data, surveillance data, host-pathogen interactions and orthology profiles across all integrated organisms." (Article - VEuPathDB NAR 2023, p. 809)

**demographics - primary data - unique**

data that could not be obtained elsewhere or recreated, often obtained directly from the researchers who created it, either through deposit or invitation to deposit

example: "We have enjoyed access to rich sources of microscopic data from several key laboratories, especially those at the MRC/LMB (Cambridge, England), the University of Wisconsin, Caltech, the University of Texas, and Johns Hopkins University. Our archival collection of TEM images in New York now includes most of the work done at MRC/LMB, including the unpublished notes of John Sulston, John White, Richard Durbin, and Donna Albertson..." (Webpage - WormAtlas WBM 2002, p. 1)

**demographics - primary use**

reported research use and rationale for the resource

example: "MorphoBank's effort has been to standardize, consolidate, organize. Just make it more consistent. FAIR acronym is one of the ways that I try to talk about adhering to all of those principles for the scientific community." (Video - MorphoBank: 195-198)

**demographics - web launch yr**

year first appeared online, specifically via the world wide web

example: "In 1995, OMIM was developed for the World Wide Web by NCBI, the National Center for Biotechnology Information." (Webpage - OMIM About, p. 1)

**distributed support - ingest workflows**

reliance on methods to obtain structured data from external resources

example: "We are utterly dependent upon people sequencing genomes, depositing those in NCBI, and NCBI curating those genomes. So we're harvesting data that other people are generating." (Video - rrnDB: 477)

**distributed support - literature**

reliance on the availability of scientific literature relevant to the mission and use of the resource

example: "Everything in OMIM is based on the peer-reviewed biomedical literature. If it's not in a paper, it's not in there." (Video - OMIM: 464)

**distributed support - volunteer service**

reliance on unpaid, volunteer labor to carry out activities important to the function of the resource example: "There's a lot of volunteer effort that comes from all of those member and affiliate organizations." (Video – BHL: 82)

**fit-for-purpose - authoritative**

provision of summaries, interpretations, and reviews created directly from SME or informed by SME (e.g., review of community-provided annotations, editorial review of submissions, etc.)

example: "Automated analyses and summaries (including experimental metadata) integrated with the above, along with annotations available from professional curators or contributed by community experts." (Webpage - VEuPathDB About, p. 1)

**fit-for-purpose - custom tools**

tools developed to aid in search/retrieval, analysis, workflow, or visualization

example: "We wrote several Open Source applications in Go and Scala to detect candidate scientific names then verify them as names by comparing them to 27 million scientific name-strings aggregated by GNA." (Article - BHL Tools 2019, p. 2)

**fit-for-purpose - expert annotation**

use of SME to enhance data records

example: "Another resource, Slidable Worm, provides an interactive and annotated series of EM images through a wild-type adult hermaphrodite. Descriptions of all 302 hermaphrodite and 385 male neurons, anatomical methods, and quick links to key anatomical publications are also included as part of the resources in WormAtlas." (Article - WormAtlas JNem 2021, p. 1)

**funding variability - as an aim**

resource receiving monetary funding as one of several outcomes of a research proposal

example: "2) to examine how and why certain biological attributes (e.g., molecules, morphology) have evolutionary histories that do not appear to match the overall family tree. We aim to make the results of our morphological work as standardized as possible with published illustrations." (Document - MorphoBank NSF 9903964, p. 3)

**funding variability - change over time**

nature of funding changed over time

example: via memo "page describes funding history, including initial grants, subsequent grants, donations, current membership, and host/member support" (Webpage - BHL Funding, p. 1)

**funding variability - dedicated for launch**

received direct monetary support to initiate the resource

example: "NIH Grants P01 DC 04732 and G08 LM05583 [a NLM Integrated Academic Information Management Systems grant] from the National Library of Medicine support this work." (Article - ORDB NAR 2002, p. 7)

**funding variability - for operations**

received monetary support that funds the operation of the resource

example: "From 2014-2024, the annual operating budget for VEuPathDB has averaged ~$8.5M in total costs; 6M in direct costs, of which ~60% has been provided by an NIAID Bioinformatics Resource Center contract, plus additional funding from the Wellcome Trust Resource Grants program and others. (Webpage - VEuPathDB FAQ, p. 2)"

**funding variability - not explicit**

no monetary support via grants/contracts received, e.g., not named as an aim and instead supported through indirect costs or other sources

example: via memo "describes multiple attempts to secure funding over years, at first including the resource as an aim and then eventually dropping it" (Video - ORDB: 228-231)

**funding variability - projects/features**

received monetary support to develop new features or conduct resource-related projects

example: "National Science Foundation (NSF) | Missouri Botanical Garden: Global Names Project, 2011 – 2014 | $295,000" (Webpage - BHL Funding, p. 1)

**host support - leadership/faculty**

reliance on host organization for support of faculty who in turn provide resource leadership and management

example: "Victor McKusick [faculty at Johns Hopkins University] has carried out the authoring of MIM more or less single handedly for the last thirty years." (Article - OMIM NAR 1994, p. 1)

**host support - misc**

additional host resources provided on an ad hoc basis, through unique opportunities, etc.

example: "We will be able to leverage key resources at Illinois to help strengthen the project. For example, part of the current project will include folks from the National Center for Supercomputing Applications Advanced Visualization Lab to develop 3D models of C. elegans anatomy." (Webpage - WormAtlas ACES Annouc 2023, p. 2)

**host support - shared services**

reliance on administrative, financial, technical, or infrastructure resources offered through the primary host organization example: "The storage is all under at the Center for Biotechnology and Genomics, which is my home base at Texas Tech." (Video - ORDB: 517-518)

**impact - current**

expressed impacts currently experienced at the time of writing in summer 2025

example: "We don't know if they're indirect costs are going to change. We don't know this, so we've got to adjust." (Video - rrnDB: 454-456)

**impact - deterioration of field**

expressed impact of declining health of the research field overall due to lack of standardization, authoritative and easy access information

example: "And then every person will have their own interpretation. Like they did before." .(Video - MorphoBank: 194-195)

**impact - efficiencies**

expressed impact of research made more difficult, less complete, and/or more time consuming

example: "Without Morphobank, it wouldn't be reproducible. We wouldn't be able to either reanalyze it or to reuse it and build on it." (Video - MorphoBank: 221-223)

**impact - lost data**

expressed impact of loss of unique primary data that could not be obtained elsewhere or recreated, especially relevant to resources obtaining otherwise unavailable (i.e., usually acquired from researchers directly)

example: "Researchers studying niche organisms like Aspergillus, Candida, and other fungi; Toxoplasma, Plasmodium, and other parasites; and arthropod disease vectors emphasize that the specificity, breadth, and depth of VEuPathDB's data cannot be easily replicated elsewhere." (Document - VEuPathDB Full Survey, p. 5)

**impact - lost opportunities**

expressed impacts of not doing or continuing work to expand, update, or innovate with the resource

example: "... browsers currently support it, but at some point, they may stop supporting it. So that's something we're actually trying to work on, and we would put into a proposal that we need to update this. Otherwise, it may not be sustainable." (Video - WormAtlas: 187-189)

**impact - lost service**

expressed impact of value of what the database does for user beyond access to the data itself, e.g., curation, annotation, aggregation, etc.

example: "Loss of resource centralization (i.e., no "one-stop shop")" (Webpage - VEuPathDB Survey Results, p. 1)

**impact - SME**

expressed impacts of losing deep professional knowledge of data, topic, and/or infrastructure built up over time

example: "... we have people who are experts and have functionally serve the BHL, some of them almost since its inception. We do not know how we can support these functions in the immediate term. And while we have to look at the big picture of how we support the BHL, these are the people with all the expertise and institutional knowledge, and we risk losing all of that." (Video - BHL: 337-338)

**prior issues - disagreements**

described as confusion over or disagreement in how data were interpreted

example: "This is critical to do at this time because scientists working on different data types continue to find different answers as to who is the closest relative of whales: living taxa indicate that it is the hippopotamus and extinct taxa indicate that it is mesonychians." (Document - MorphoBank NSF Grant, p. 3)

**technical - API**

development of an API to access data

example: "The MorphoBank public API supports HTTP GET requests for published resources. API documentation is available here." (Webpage - MorphoBank FAQ, p. 23)

**technical - cloud**

migration to cloud storage

example: via memo "notes that the infrastructure has migrated to AWS" (Interview Notes - DB1)

**technical - code base**

substantial revision of all or part of the code base to update infrastructure, expand features, etc.

example: "EuPathDB has undergone dramatic changes specifically designed to enhance data accessibility and provide database users with a graphical query-building interface that effectively creates a venue for constructing complex search strategies with relative ease." (Article - VEuPathDB NAR 2010, p. 2)

**technical - migration**

specifically moving technical infrastructure from one host organization to another

example: "The Smithsonian is not going to be able to host the secretariat or the infrastructure anymore." (Video - BHL: 191)

**technical - scrapping**

reported bot use scrapping data resource websites

example: "We are all so bombarded, all the databases are being screen-scraped constantly at very high volume." (Video - OMIM: 519)
